# Fear of parasitism affects the functional role of ecosystem engineers

**DOI:** 10.1101/2021.08.27.457894

**Authors:** Kim N. Mouritsen, Nina P. Dalsgaard, Sarah B. Flensburg, Josefine C. Madsen, Christian Selbach

## Abstract

Fear is an integrated part of predator-prey interactions with cascading effects on the structure and function of ecosystems. Fear of parasitism holds a similar ecological potential but our understanding of the underlying mechanisms in host-parasite interactions is limited by lack of empirical examples. Here, we experimentally test if blue mussels *Mytilus edulis* respond behaviourally to the mere presence of infective transmission stages of the trematode *Himasthla elongata* by ceasing filtration activity, thereby avoiding infection. Our results show that blue mussels reduced clearance rates by more than 30% in presences of parasites. The reduced filtration activity resulted in lower infection rates in experimental mussels. The identified parasite-specific avoidance behaviour can be expected to play a significant role in regulating the ecosystem engineering function of blue mussels in coastal habitats.

## Introduction

Identifying factors and processes that determine the structure of natural communities, and in turn ecosystem functioning, continues to be a main goal in ecological research. Across ecosystems, a very narrow subset of organisms is often found to play disproportionally great ecological roles, and therefore function as designated ecosystem engineers or keystone species (Jones et al. 1994, Power et al. 1996, Jones et al. 1997, Bertness 2007). The blue mussel *Mytilus edulis* is such an ecological important species in North Atlantic inter- and subtidal communities. They form extensive mussel beds along the coastline where they fulfil central ecological roles, such as filtering out and biodepositing organic matter, and creating biogenic reefs on which other organisms either depend for shelter, substrate or foraging, or are totally excluded as inferior space competitors (Ragnarsson and Raffaelli 1999; Bertness 2007; Norling and Kautsky 2007, Commito et al. 2008). Consequently, any organism that influences the functional performance of blue mussels may itself be a significant player in the coastal ecosystem. A prominent example of such an indirect ecological role is predation from sea stars on intertidal mussels. If sufficiently abundant, sea stars may reduce mussel densities to levels that allow establishment of otherwise excluded species of sessile organisms, thereby elevating local biodiversity (e.g. Pain 1980, Kaiser et al. 2020).

Not only predators may play such indirect ecological roles, though. Blue mussels are commonly host to a range of parasites with significant pathological consequences. Among those, the ubiquitous trematode *Himasthla elongata* utilizes blue mussels as an intermediate host that becomes infected by free-swimming parasite stages (cercariae) emitted from the common periwinkle snail *Littorina littorea* (Werding 1969). Once inside the mussel, the parasites encyst as metacercariae in the tissues and can reduce the hosts attachment to the substrate, reduce growth rates as well as render the mussel more vulnerable to predation by definitive shorebird hosts and other bivalve-eating predators (Lauckner 1983, Beck 2008, Bakhmet et al. 2017).

Aside from these obvious direct consumptive effects of infection, parasites like *H. elongata* can also induce more cryptic non-consumptive effects with marked ecological ramifications. Comparable to the ‘ecology of fear’ described for predator-prey interactions, the fear of parasitism has been shown to play a significant role in various host-parasite systems (Rohr et al. 2009, Marino and Werner 2013, Buck et al. 2018, Buck 2019). However, fear-driven parasite avoidance behaviour remains particularly understudied in aquatic environments, and its ecological impact difficult to assess (Behringer et al. 2018). Recent findings indicate that blue mussels attempt to avoid the detrimental *H. elongata* infections by closing their siphons, through which infective cercariae mainly enter (Selbach and Mouritsen 2020). Such a parasite avoidance strategy may be costly to the mussel, as it prevents filter-feeding when parasites are present and could have drastic implications for the bivalves’ ecosystem engineering potential (Selbach and Mouritsen 2020). However, to what extent this fear of parasitism actually controls the filtration activity of blue mussels is currently unknown and remains to be investigated. Here, we test this intriguing hypothesis in a series of microcosm experiments and predict (1) that the mussels’ filtration activity is significantly reduced in presence of *H. elongata* cercariae in the water column, (2) that this response is parasite-specific and not elicited by presence of inert particles, and (3) that this parasite-avoidance behaviour results in lower trematode infection rates in mussels. Confirmation of our working hypotheses will have important implications for the ecosystem engineering role of blue mussels, as individuals ceasing filtration due to fear of parasitism will remove less organic matter from the water column, deposit less faeces and likely experience lower growth rates.

## Material and methods

### Collection of animals

All animals were collected ultimo June 2020, a few days prior to experimentation. Blue mussels (*Mytilus edulis*) were obtained from mariculture (Danish Shellfish Centre, Mors, 56.7879N/8.8778E) and therefore free of trematode infections (Buck et al. 2005, Selbach et al. 2020). The mussels were sorted according to size, and individuals of c. 35 mm in maximum shell length were retained and stored in running filtered seawater (c. 18°C, 25 PSU) until experimentation. Common periwinkles (*Littorina littorea*) harbouring the infective larval stages (cercariae) of the trematode *Himasthla elongata*, were collected in Knebel Bay (56.2086N/10.4796E) and stored dry at 18°C until extraction of parasite larvae a few days later.

### Experimental design

To determine the relative impact of swimming *H. elongata* cercariae and suspended inert microplastic particles on the filtration activity of blue mussels, we performed filtration experiments with four unique experimental treatments: (1) addition of parasites only, (2) addition of microplastic only, (3) addition of both parasites and microplastic, and (4) control, i.e. absence of both parasites and microplastic. Including microplastic as an experimental treatment allowed us to clarify whether mussels react specifically upon parasites or any suspended particle touching the mussel tissue.

The experimental unit was a 2L PP container (Ø = 13.3 cm) containing 1.5L solution of microalgae serving as food for the experimental mussels. The solution was made from pre-filtered seawater (<5 μm, 25 PSU) and added concentrated paste of *Tetraselmis* sp. cells (10-12 μm, Tetraselmis 3600™ from Reed Mariculture), resulting in a final concentration of 20 mio cells L^−1^. Each unit, planned to enclose a single experimental blue mussel, was equipped with an efficient air supply to ensure full oxygen saturation as well as ample water circulation. A total of 20 such experimental units were placed on a table in a temperature-controlled chamber (18°C), of which five received each of the four experimental treatments. Additional three units were established to account for the sedimentation rate of algae during the experiment, and hence, these three units did not receive any of the four treatments, nor any mussels. The experimental units were placed haphazardly on the table according to received experimental treatment.

At commencement of the experiment, parasite-treated units each received 200 recently released *H. elongata* cercariae (≤ 3 hours of age). After emergence from the snail first intermediate host, these parasite larvae curl-up and use beating tail movements to stay buoyant and for propulsion through the water column. The spherical body of the swimming cercaria has a diameter of 250-300 μm. The microplastic-treated units each received c. 1500 white PVC particles (Ø = 150 μm), kept in solution by the currents created by the submerged air bubbler. The somewhat smaller PVC particle size in comparison to the cercariae was necessary in order to keep the majority in solution at the applied level of water circulation. As a trade-off, a greater number of PVC particles than cercariae was added to each experimental unit.

Release of cercariae from host snails was induced by incubating the snails for 2 hours in seawater-filled glass jars under illumination at c. 25°C. The swimming parasites larvae was then extracted and enumerated by pipetting them into petri dishes, and from there poured (c. 5 mL) into the designated experimental units. The 1500 PVC particles planned for each microplastic-treated unit were administered as weight equivalents dissolved in 5 mL seawater and then poured into the experimental units. Units not receiving any of these treatments were added 5 mL of pure filtered seawater.

### Experimental protocol

Prior to experimentation, a single mussel individual, haphazardly chosen from the storage tank, was placed in each experimental container together with one litre of filtered, algae-free seawater and air-supply for a three-hour acclimation period. During this period, the mussel attached itself to the bottom of the container by its byssus threads. Following acclimation, the filtered seawater was replaced by a similar amount (1L) of the prepared algae solution (see above) after which each experimental container received the pre-designated treatment. Additional algae solution was then added in order to reach a final water volume of 1.5L in all containers.

Following two hours of experimentation, during which time it was ensured that all mussels were alive and filtering (i.e. showing emerged, active siphons), a 0.5L water sample was taken from each experimental unit and analysed for chlorophyll-a content. The experimental mussels were individually stored dry in glass jars for 12 hours to allow transmitted parasite larvae to encyst as metacercariae in the host tissue. Mussels were then transferred to plastic bags and frozen until dissection.

Because of the labour involved in obtaining sufficient numbers of parasite larvae of required age as well as in processing collected water samples post-experimentally, five replicas per treatment plus three sedimentation controls were considered the limit for one experimental run (see design above). Hence, to increase statistical power the experiment was repeated on three consecutive days, resulting in a total of 15 replicas per treatment and nine sedimentation controls. However, this also meant that day was introduced as an additional factor in the experimental design.

### Chlorophyll-a measurement and mussel dissection

The 0.5L water samples from each experimental unit were filtered individually through a tissue filter (Advantec GC-50® glass fiber filter, retention: 0.5 μm) using a vacuum pump. Filters containing retained microalgae were transferred to dark test tubes together with 5 mL 96% ethanol and stored dark for 12 hours. The samples were then placed in an ultrasonic cleaner for 5 min followed by 20 sec on a shaker and finally 10 min centrifugation (6000 rpm). The resulting supernatant of each sample was transferred to a spectrophotometer (Spectronic Helios Alpha®) and the absorbance was measured at 750 and 665 nm. Subsequently, the chlorophyll-a concentration (μg L^−1^) was calculated according to Riemann et al. (1989).

The experimental mussels were dissected and screened for the presence of metacercariae and microplastic in their tissues. Prior to dissection, the mussels’ shell length was measured accurately (nearest 0.1 mm) using an electronic calliper. Subsequently, the soft tissue was removed and squeezed between two glass plates and viewed under a stereomicroscope for enumeration of metacercariae and plastic particles present in the various tissues (foot, gills/palps, mantle/siphons, digestive tract, remaining visceral mass).

### Data analyses

Clearance rate (*CR*, L hour^−1^) was used as an estimate of the mussels’ filtration activity and targeted as dependent variable in the analysis. *CR* = (*V*/*t*) * ln(*C_0_*/*C_t_*), where *V* = water volume (L), *t* = experimental period (hours), *C_0_* = chlorophyll-a concentration at *t* = 0 (μg L^−1^) and *C_t_* = chlorophyll-a concentration at the end of the experimental period (e.g. Coughlan 1969). Prior to analysis, *CR* values were corrected for sedimentation of suspended microalgae on the three experimental days.

The statistical analysis was carried out in IBM SPSS 26. ANOVAs were preceded by tests of homogeneity of error variance (Levene’s test) and evaluation of normality, and the presented reduced models were preceded by full models demonstrating absence of factor interactions. Data from five experimental units were excluded in the analyses. Three were excluded because the mussels spawned during the experiment, affecting the reliability of the spectrophotometric measurements, and two were lost because of apparatus error. Hence the effective sample size per treatment per day ranged between 3 and 5. Standard error (SE) is given throughout as variation around the central tendencies.

## Results

### Mussel size

The shell lengths of the experimental blue mussels were statistically very similar across treatments and experimental days (Electronic supplement, Table S1). Hence, mussels size as a potential confounding factor was omitted in further analyses. The grand mean shell length of applied mussels was 34.5 ± 1.5 (SE) mm (N=60).

### Impacts on clearance rate

The mussels’ clearance rate was significantly affected by parasite exposure and experimental day, respectively, explaining 26 and 28% of the variation, whereas presence of microplastic particles had no bearing on filtration activity at all (Table 1). No statistically significant interactions were evident among the three independent factors (P ≥ 0.188; 0.188-0.824). Hence, presence of parasite larvae in the surrounding water generally decreased the clearance rate by 32.1% compared to absence of parasites, and clearance rates increased more or less steadily with time (experimental day) regardless of the treatment (Fig. 1).

**Table 1.** Summary statistics of reduced model 3-way ANOVA including clearance rate (L hour^−1^) of *Mytilus edulis* as dependent variable and parasite exposure (presence/absence of *Himasthla elongata* cercariae), microplastic exposure (presence/absence) and experimental day (1-3) as fixed factors. Partial η^2^ denotes effect size, i.e. the proportion of variance explained. Preceding full model ANOVA showed absence of significant 2- and 3-way interactions (P = [0.188; 0.824]).

| Source | df | F | P | Partial $\eta^2$ |
| --- | --- | --- | --- | --- |
| Parasites | 1 | 17.942 | <0.0005 | 0.264 |
| Microplastic | 1 | 0.343 | 0.561 | 0.007 |
| Day | 2 | 9.595 | <0.0005 | 0.277 |
| Error | 50 |  |  |  |

**Figure 1.**
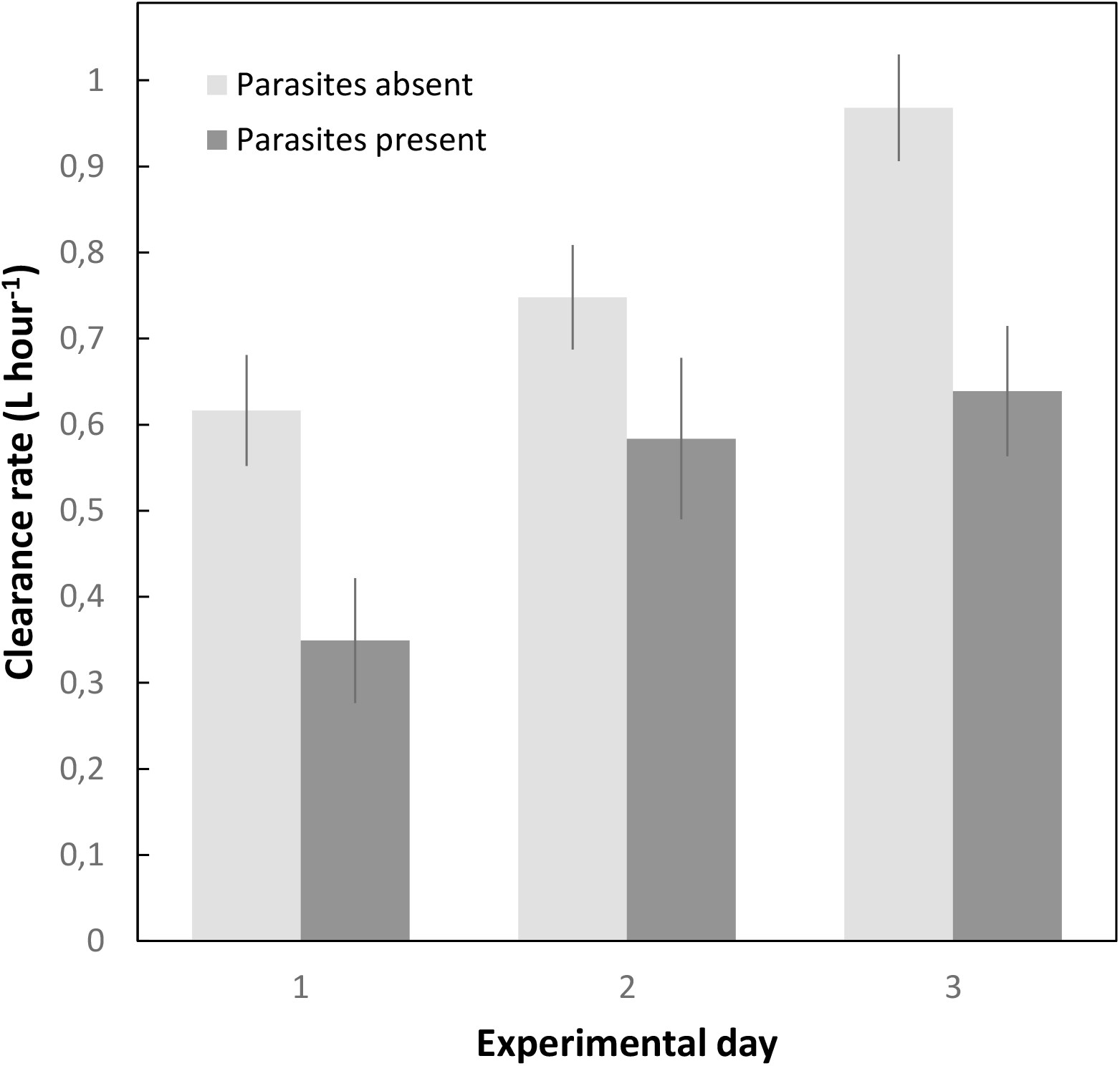
Clearance rate (mean L hour^−1^ ± SE, n = 8-10) of experimental blue mussels *Mytilus edulis* in presence and absence of infective parasite stages (*Himasthla elongata* cercariae) as a function of experimental day. See Table 1 for summary statistics.

### Within-mussel distribution of parasites and microplastic

All mussels exposed to parasites became infected during the experiment and the overall infection success was 10.8 ± 1.6 (SE) % (i.e. averagely >20 encysted metacercariae mussel^−1^). The parasites were not evenly distributed across the different tissues: the main site of infection was the foot but also mantle/siphons and gills were targeted (Table 2). Particles of microplastic were found in connection to all types of tissues but particularly on the surface of mantle, siphons and the visceral mass. A major proportion was also located in the digestive tract and hence directly ingested by the mussels (Table 2). On average, we recovered 14.9 ± 5.3 (SE) PVC particles per exposed mussel.

**Table 2.** The distribution (%) of *Himasthla elongata* metacercariae and microplastic particles in and on various tissues of exposed experimental blue mussels *Mytilus edulis*. Summary statistics from Goodness-of-fit tests analysing even distribution across tissues are given.

|  | Proportion (%) |  |  |  |  |  |  |  |
| --- | --- | --- | --- | --- | --- | --- | --- | --- |
| Parameter | Foot | Mantle and siphons | Gills and palps | Digestive tract | Other tissues | N | X<sub>4</sub><sup>2</sup> | P |
| Metacercariae | 65.9 | 15.9 | 17.7 | 0 | 0.5 | 646 | 623.9 | <0.0005 |
| Microplastic | 0.2 | 22.9 | 0.9 | 36.5 | 39.5 | 446 | 316.0 | <0.0005 |

### Infection success and clearance rate

The infection success of *H. elongata* during the 2-hour experimental period was independent of microplastic treatment as well as experimental day (Electronic supplement, Table S2). Hence, after combining all data, a significant positive relationship between infection success and mussel clearance rate could be established (Fig. 2). Although the residual variation is large, this suggests that a major route of entry by the parasites indeed goes via the filtration current of the host.

**Figure 2.**
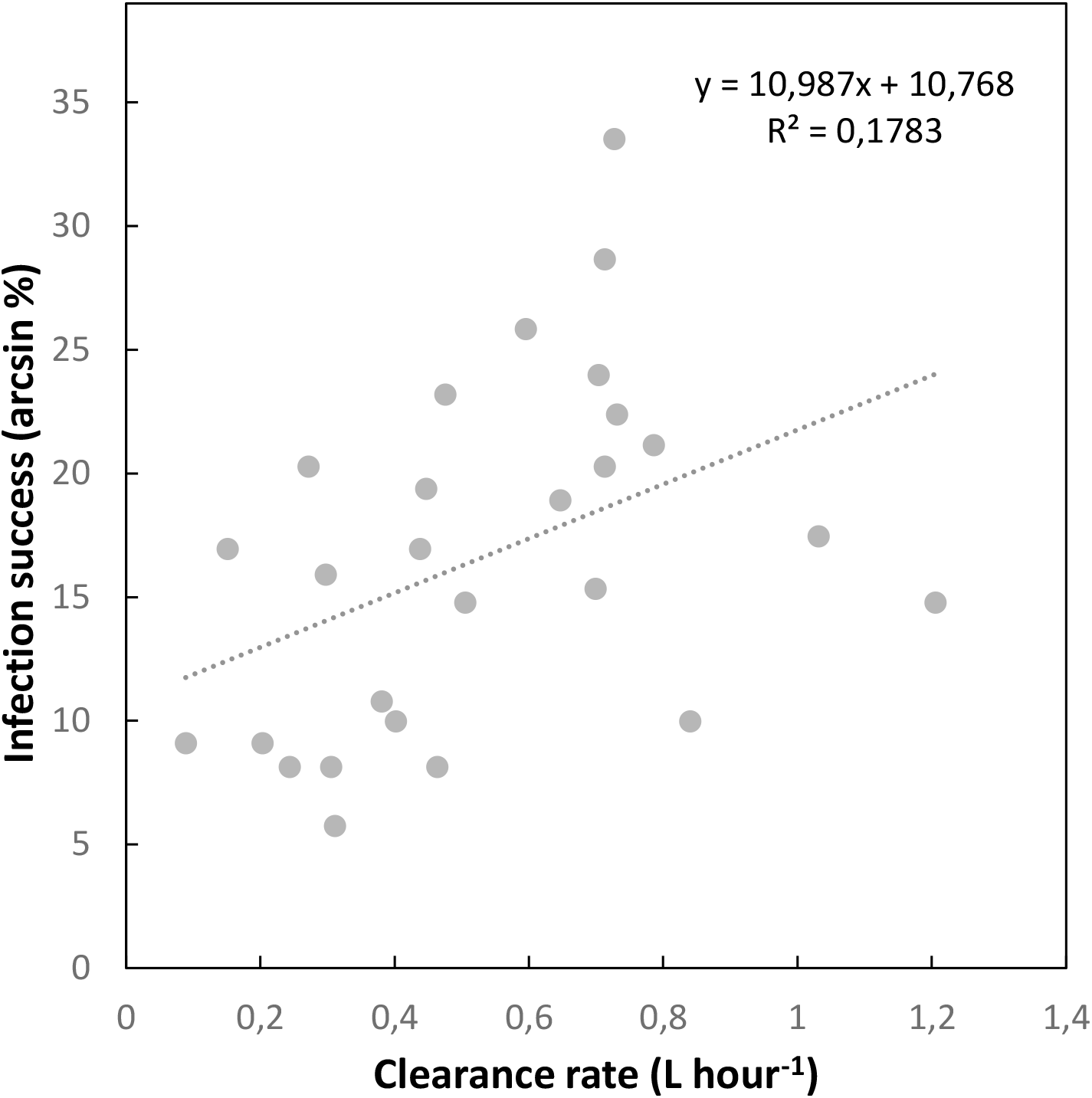
Infection success (arcsin %) of experimentally added *Himasthla elongata* cercariae as a function of the clearance rate (L hour^−1^) of host mussels (*Mytilus edulis*). Linear regression, r^2^_25_ = 0.178, P = 0.028.

## Discussion

In this study, we tested if the ‘fear’ of parasites can induce behavioural changes in blue mussels *Mytilus edulis* and potentially affect the ecosystem engineering role of these abundant bivalves in coastal systems. The present results provide unequivocal evidence that the mere presence of infective *Himasthla elongata* cercariae in the immediate surroundings elicits a significant reduction in blue mussel filtration activity. Quantitatively, clearance rates were reduced by more than 30%, which is a surprisingly high number comparable to the pathological impact recorded in mussels heavily infected by trematodes directly targeting their feeding apparatus (Stier et al. 2015). Moreover, a reduced clearance rate resulted in lower trematode infection rates, highlighting the effectiveness of this parasite-avoidance behaviour.

In the present case, the mechanism is rather a behavioural response by the mussel (siphon closing), most likely induced by mechanical contact between swimming parasite larvae and the mussels exposed mantle and siphon tissue during filtration. Aside from being in accordance with direct observations of host-parasite interactions under laboratory conditions (Selbach and Mouritsen 2020), the alternative possibility of a pathological effect can readily be excluded by the very short experimental period (two hours) and an independent line of evidence demonstrating absence of any impact of mature *H. elongata* infections on host filtration (Selbach et al. in prep.).

Interestingly, the mussels’ siphon closing appears induced specifically by presence of parasites. Inert particles in the water column, on the other hand, do not elicit a similar response, evidenced by similar clearance rates in absence and presence of suspended microplastic (Table 1). Logically, living in a highly dynamic coastal habitat with frequent wave- or tide-induced resuspension of sediment particles, mussels cannot afford to cease feeding whenever a small foreign item touches its tissues. Accordingly, the experimentally added plastic particles were found throughout the mussels’ mantle cavity and even in the intestine, evidencing their direct consumption (Table 2). Although microplastic particles have been shown to reduce clearance rates in *Mytilus* spp., these reductions in filtration result from histopathological effects following particle ingestion rather than uptake avoidance behaviour (Woods et al. 2018, Alnajar et al. 2021). Hence, via mechanoreceptors present on the tissue surfaces blue mussels seem capable of recognizing and distinguishing the specific pattern of touch created by cercariae and in turn launch the necessary defence measures to avoid their entrance by retracting their siphons. Additionally, bivalves have been shown to respond to chemical risk cues from predators or alarm signals from conspecifics (Kobak and Ryńska 2014, Dzierżyńska-Białończyk et al. 2019). It therefore remains to be tested, if blue mussels can similarly react to the presence of chemical cues of infective trematode cercariae in the environment.

Blue mussels have all reasons to fear and avoid these parasites. Although metacercariae of *H. elongata* can be found in most host tissues, cercariae specifically target the muscular foot of the mussel (Table 2, Lauckner 1983). Here, they immobilize the foot, reducing its capability to deposit byssus threads necessary for anchoring the mussel to the substrate. Consequently, heavily infected mussels become easy prey to shorebird definitive hosts as well as other mussel-eating predators (Lauckner 1983, Beck 2008). The increased mortality rate caused by such parasite-induced facilitation of trophic transmission (parasitic manipulation) has not yet been fully quantified in the *Mytilus-Himasthla* system. However, in a comparable host-parasite system the predation rate on parasite-manipulated bivalves was found to be 5-7 fold higher than in unaffected specimens (Thomas and Poulin 1998, Mouritsen 2004). This emphasizes that the selection for effective anti-parasite measures in bivalves is strong, and the cost of parasite infections likely outweighs the costs of parasite avoidance (Poulin et al. 1999).

This pressure likely drives the evolution of fitness-improving adaptive measures to avoid infection by trematode stages, including perception abilities to distinguish between parasites and benign particles as well as an effective way of avoiding the parasites entering the mussels. Regarding the latter, an important route of infection follows the mussels’ filtration current, evidenced by the positive relationship between infection success and clearance rates of the experimental mussels (Fig. 2). Hence, ceased filtration activity appears the obvious and most effective way to prevent parasites from entering the mantle cavity and from there penetrating the mussel’s tissues. Since other parasite species that occur in sympatry, such as the trematode *Renicola roscovita*, infect blue mussels via the same infection pathways, potential interactions of non-consumptive effects of different parasites are likely drivers of co-evolutionary dynamics of this host-parasite system (Thieltges and Rick 2006).

Clearance rates increased steadily during the course of the 3-day experiment (Fig. 1, Table 1). As mussels were stored in filtered, and hence food deprived, seawater prior to experimentation, the most parsimonious explanation for this temporal pattern is increasing hunger motivating mussels to feed. It is therefore intriguing that there was no significant day-parasite interaction evident in measured clearance rates (Table 1). This means that the negative impact of parasite presence on clearance rates decreased with time (see Fig. 1), which in turn points at a trade-off between hunger and fear of parasites: a hungry mussel is willing to take a greater risk of infection. Similar state-dependent trade-offs between food acquisition and risk avoidance have been demonstrated for various predator-prey interactions (Juliano et al. 1993, Pettersson and Brönmark 1993), although these patterns do not seem universal (see Horat and Semlitsch 1994). The state- or context-dependency of parasite avoidance dynamics should therefore be tested for different host-parasite systems.

This non-consumptive parasite effect, driven by the mussel hosts’ attempt to reduce their parasite load, can be expected to have far-reaching effects on the coastal ecosystem. As a well-defined ecosystem engineer, blue mussels generally acts as a dominant space competitor excluding most macrophytes and other sessile macrofaunal organisms from the mussel-bed (Pain 1980, Kaiser et al. 2020). On the other hand, their great filtration capacity establishes a significant benthic-pelagic coupling where organic matter from the water column is deposited on the seabed as faeces and pseudofaeces. This, in turn, facilitates a rich community of smaller in- and epifaunal organisms. In addition, mussel beds usually attract a diverse guild of mussel predators, such as crabs, sea stars and shorebirds. Hence, a marked reduction of the mussels’ feeding activity due to the mere presences of parasites will immediately reduce the transport of organic material into the substrate thereby inhibiting the benthic animal community. On a longer time-scale, reduced feeding rates will translate into reduced growth rates, to the benefit of the predator guild that will find a greater proportion of mussels below the size refuge. Parasite-induced fear is therefore likely to act as a major structuring force in intertidal communities.

These potential ecological ramifications, however, rest on the assumption that the parasite-mediated decrease in filtration activity is not off-set by higher activities at times where parasites are less abundant in the water column (see Selbach and Mouritsen 2020), or when increasing hunger levels override the parasite-avoidance behaviour of filter feeders (see above). However, considering the wide-spread occurrence of the focal parasites, by which 20-40% of the first intermediate snail host population typically is infected (e.g. Werding 1969, Mouritsen and Halvorsen 2015, Mouritsen 2017), there is good reason to expect an ecological impact of the identified non-consumptive parasite effect also *in situ*. As previously stressed (Selbach and Mouritsen 2020), such an impact might well surpass the direct consumptive effects of trematode infection in blue mussels. Hence, further unravelling how fear of parasitism influences host behaviour and thereby shapes ecosystems will be a decisive step towards a better understanding of how coastal ecosystems are organised and functioning.

## Supporting information

Original Data S3

## Data archiving statement

The datasets supporting this article have been uploaded as part of the supplementary material (Electronic supplement S3).

## Ethics statement

All institutional and national regulations for the care and use of animals were followed.

## Acknowledgements

We are grateful to the Danish Shellfish Centre, Mors, Denmark for providing us with blue mussels for the experiment We thank Jessica Schwelm for feedback on an earlier draft of the manuscript.

