## Supplemental Table S1 and S2 for "Fear of parasitism affects the functional role of ecosystem engineers"

**Electronic supplement**

Journal: Preprint

Year: 2021

**Table S1.** Summary statistics of full model 2-way ANOVA including shell length (mm) of experimental blue mussels *Mytilus edulis* as dependent variable and treatment (presence/absence of parasite and microplastic) and experimental day (1-3) as fixed factors. Grand mean: 34.53 ± 1.45 mm (n = 60).

| **Source** | **df** | **F** | **P** |
|---|---|---|---|
| Treatment | 3 | 1.152 | 0.676 |
| Day | 2 | 0.416 | 0.662 |
| Treatment x Day | 6 | 0.848 | 0.540 |
| Error | | | |

**Table S2.** Summary statistics of reduced model 2-way ANOVA including infection success of *Himasthla elongata* (arcsin-transformed %) in experimental *Mytilus edulis* individuals as dependent variable and microplastic exposure (presence/absence) and experimental day (1-3) as fixed factors. Partial $\eta^2$ denotes effect size, i.e. the proportion of variance explained.

| **Source** | **df** | **F** | **P** | **Partial $\eta^2$** |
|---|---|---|---|---|
| Microplastic | 1 | 0.681 | 0.417 | 0.026 |
| Day | 2 | 0.523 | 0.599 | 0.039 |
| Error | 26 | | | |
